# Molecular detection using hybridization capture and next-generation sequencing reveals cross-species transmission of feline coronavirus type-1 between a domestic cat and a captive wild felid

**DOI:** 10.1101/2024.01.02.573944

**Authors:** Ximena A. Olarte-Castillo, Laura B. Goodman, Gary R Whittaker

## Abstract

Feline coronavirus (FCoV) infection normally causes mild or subclinical signs and is common in domestic cats. However, in some cats, FCoV infection can also lead to the development of feline infectious peritonitis (FIP)—a typically lethal disease. FCoV has two serotypes or genotypes, FCoV-1 and FCoV-2, both of which can cause FIP. The main difference between the genotypes is the viral spike (S) protein that determines tropism and pathogenicity, crucial mechanisms in the development of FIP. Subclinical infection and FIP have both been reported in wild felids, including in threatened species. Due to the high genetic variability of the S gene and the technical challenges to sequencing it, detection and characterization of FCoV in wild felids have mainly centered on other more conserved genes. Therefore, the genotype causing FIP in most wild felids remains unknown. Here, we report a retrospective molecular epidemiological investigation of FCoV in a zoological institution in the United States. In 2008, a domestic cat (*Felis catus*) and a Pallas’ cat (*Otocolobus manul*) sharing the same room succumbed to FIP. Using hybridization capture and next-generation sequencing, we detected and sequenced the complete 3’-end of the viral genome (∼8.2 Kb), including the S gene, from both felids. Our data show for the first time that FCoV-1 can be transmitted between domestic and wild felids and extends the known host range of FCoV-1. Our findings highlight the importance of identifying the genotype causing FIP, to develop effective control measures.

## Introduction

Feline coronavirus (*Alphacoronavirus* genus, *Coronaviridae* family) is a widespread virus that is highly prevalent in domestic cats worldwide. Feline coronavirus (FCoV) infection can cause subclinical disease or mild signs of gastroenteritis. However, in a subset of cats, FCoV can cause a lethal immune-mediated disease known as feline infectious peritonitis (FIP). Due to its multi-systemic nature, FIP can have a variety of clinical signs, including neurological (1) and upper respiratory signs (2). To date, an effective vaccine to prevent FIP is not available, and while treatment with anti-viral drugs such as GS-441524 has proven successful (3), their use has not been approved in the U.S. Current evidence suggests that certain mutations in the sub-clinical biotype of FCoV (feline enteric coronavirus, FECV (4)), allow the virus to shift its tropism from epithelial cells to macrophages and monocytes, an essential step for the development of FIP (5). The mutant pathogenic biotype is known as feline infectious peritonitis coronavirus (FIPV). Identifying mutations associated with the FIP phenotype requires the characterization of the genetic variation between FCoV variants from healthy and FIP-cats.

FCoV belongs to the *Alphacoronavirus-1* species (*Alphacoronavirus* genus, *Coronaviridae* family) together with canine coronavirus (CCoV), transmissible gastroenteritis virus (TGEV) and porcine respiratory coronavirus (PRCV) (6). The genome of these coronaviruses (CoVs) is around 32 kb long and includes non-structural (replicase 1a and b), structural (spike, matrix, envelope, nucleocapsid), and accessory genes (Figure 1). FCoV has two serotypes or genotypes: Type 1 (FCoV-1) and Type 2 (FCoV-2), both of which can cause FIP (7) (8). FCoV-2 is a recombinant genotype of FCoV-1 that acquired its spike (S) gene from CCoV type 2 (CCoV-2) through double homologous recombination (9) (Figure 1). The S protein contains the major antigenic sites of CoVs and is also involved in essential processes for host cell entry (10), including receptor binding (11) and fusion (12), which occur through the S1 and S2 domains, respectively. Therefore, viral molecular mechanisms related to immune evasion, viral tropism, pathogenicity, and host range differ between FCoV-1 and FCoV-2 because their S protein is highly divergent (<54% pairwise amino acid identity, (13)). For example, FCoV-2 can easily grow in cell culture, and its receptor (aminopeptidase N, APN) and receptor binding domain (RBD) have been identified and characterized (11) (14) (15) (Figure 1). In contrast, FCoV-1 is difficult to grow in cell culture, does not use APN as a receptor (16) (17), and to date, its receptor has not been identified. Consequently, to understand the mechanisms of pathogenicity of FCoV that result in FIP, it is essential to study the genetic diversity of the S protein of both FCoV-1 and -2.

**Figure 1.**
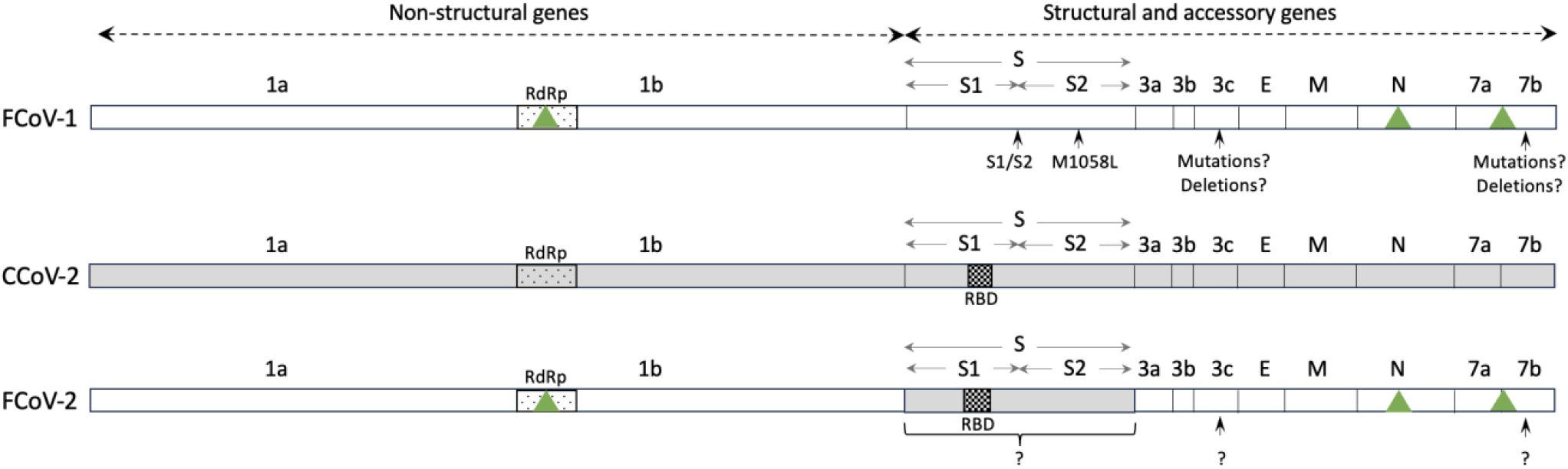
Schematic representation of the genomes of FCoV-1, CCoV-2, and FCoV-2. The genome of FCoV-1 is shown in white, and the genome of CCoV-2 in gray. FCoV-2 is a recombinant genotype that has the S gene of CCoV-2 (in gray) and the rest of the genome of FCoV-1 (in white). The overall percentage of nucleotide similarity between the S gene of FCoV-1 and CCoV-2 is <54%. The genomes are divided into non-structural (replicases 1a and 1b), structural (S, E, M, N), and accessory (3a, b, c, 7a, b) genes. The names of each gene are displayed above each genome representation. Whitin replicase 1b is the RNA-dependent RNA polymerase (RdRp) gene (displayed as a dotted box on each genome). Within the S gene of FCoV-2 and CCoV-2, the receptor binding domain (RBD, shown as a box with black squares) has been identified. The RBD of FCoV-1 has not been identified. Green triangles indicate the genes that are commonly used for the detection of FCoV in clinical samples of wild felids (RdRp, N, and 7ab). Regions in which mutations or deletions have been linked to pathogenicity in FCoV-1 and -2 are indicated by arrows in each genome scheme. Very few sequences of FCoV-2 are currently available, thus the role of some of these regions is less known (represented as question marks).

One of the major differences between the S proteins of FCoV-1 and FCoV-2 is that FCoV-1 has a so-called furin cleavage site (FCS) in the region between the S1 and S2 domains (S1/S2 region (12)). A furin-mediated cleavage in the FCS is thought to be essential for the activation of the S protein and the subsequent host receptor binding (18). Mutations and/or insertions or deletions in and around the FCS can alter the pathogenicity of FCoV by modifying the furin cleavability of the S protein (12). FCoV-1 detected in healthy cats (i.e. FECV) typically exhibit the core furin cleavage motif S(P6)-R(P5)-R(P4)-S/A(P3)-R(P2)-R(P1)|S(P1’) (in which A is Alanine, S is Serine, R is Arginine, in parenthesis the P followed by a number indicates the position of each residue within the cleavage site, and | indicates the site in which the cleavage occurs, (12)). On the other hand, mutations in this motif are typically observed in FCoV-1 recovered from cats diagnosed with FIP (i.e., FIPV, (12)). Using molecular evolutionary genetic statistical techniques, a recent study (19) reported that certain mutations in residue P4 are associated with FIPV (R in FECV; R, K, G, Q, or S in FIPV, in which K is lysine, G is glycine, and Q is glutamine). Another residue within the S protein, but outside the FCS was also detected to be relevant for the FIP-phenotype. This residue is known as residue 1058 and is methionine (M) in FECV and mostly leucine (L) in FIPV (19) (20). In comparison, although much is known about the molecular interactions between the S protein of FCoV-2 and its receptor (15), knowledge of mutations associated with pathogenicity for this genotype remains limited. Outside the S protein of both genotypes, deletions, and mutations in accessory genes 3c and 7ab have been reported in some FIPV variants (21, 22), but their exact relation with the FIP phenotype is not well established (19).

Infectious diseases can have a significant impact on wildlife populations causing substantial population declines (23), and even local extinction (24). From a conservation point of view, knowledge of the susceptibility of wild species to certain infectious diseases helps identify the best management approaches to diminish their impact on threatened populations (25). The Felidae is a diverse family within the Carnivora order that includes 38 recognized species (26), including the domestic cat (*Felis catus*). According to the International Union for Conservation of Nature (IUCN), 34% of felids (13 species) are categorized as vulnerable, 13% (5 species) as endangered, and 71% (27 species) have decreasing population trends (27). Wild felids may be susceptible to infectious diseases from domestic cats due to their close genetic relatedness. Therefore, it is important to study if common viruses of domestic cats can infect wild felids and assess their potential impact on threatened populations (28). FCoV infection has been reported in several free-ranging wild felids including endangered species like the Siberian tiger (*Panthera tigris altaica*, (29)). FIP cases have been reported from free-ranging and captive wild felids including a Mountain lion (*Puma concolor*) in California, U.S. (30), European wildcats (*Felis silvestris)* (31), and three sand cats *(Felis margarita)* (32) in zoological institutions in the UK and the U.S. The most notable case of FIP in wild felids was reported in a captive population of cheetahs (*Acinonyx jubatus*) in a safari park in Oregon, U.S. (33). After the introduction of two FCoV-infected cheetahs to the park in 1982, the virus rapidly spread infecting 100% of the cheetah population (n=45). After 4 years of the introduction of FCoV, 60% of the cheetahs died of FIP (34) (35). This example shows the impact that FCoV can have on wild felid populations and highlights the importance of assessing the prevalence of FCoV in wild species.

The S gene of both FCoV-1 and -2 has regions of high genetic variability; thus, targeting and sequencing it to detect and study the molecular epidemiology of FCoV is difficult. For example, obtaining complete or even partial sequences of S using classical techniques like RT-PCR and Sanger sequencing has proven to be challenging (36). Detection of FCoV in wild felids (free-ranging or captive) has been based mostly on serology, RT-PCR, or histology targeting more conserved genes (37). For example, commonly used serology tests target the N protein (31), and detection by RT-PCR targets genes like the RNA-dependent RNA polymerase (RdRp, (35) or 7a and 7b (38) (Figure 1). Histological detection like immunohistochemistry (IHC) and *in-situ* hybridization target the N protein (*i*.*e*., antibody FIPV3-70) and RdRp (39), respectively. While targeting these genes can be useful for detection purposes (*i*.*e*., they are highly conserved), they are not optimal for differentiating FCoV-1 and -2 (Figure 1). Therefore, although it has been reported that FCoV can be highly prevalent in certain wild felid populations (29), the prevalence of each genotype (FCoV-1 and FCoV-2) in wild felids and the specific genotype causing FIP in most previous cases is currently unknown because the S protein has not been studied. To date, two partial sequences of the RdRp gene from two cheetahs from the 1982 Oregon case (35) and a complete genome sequence of one captive sand cat (32), are the only sequences available for FCoV recovered from wild felids. Without sequence data from the FCoV genotypes circulating in wild felids, it is not possible to assess if there has been cross-species infection between domestic and wild felids. From a conservation point of view, the identification of the specific genotype that causes FIP is essential to identify the best management approaches to diminish its impact on free-ranging and captive felid populations.

The Pallas’ cat (*Otocolobus manul*) is a small felid that inhabits the steppes of Central and Western Asia. Due to their solitary lifestyle and remote habitat, little is known about the infectious diseases that affect this species (40). Increasing fragmentation of the Pallas’ cat habitat and the presence of free-ranging domestic cats from villages or those accompanying herdsmen may increase their exposure to pathogens of domestic cats (41). For example, serology assays have shown that both Pallas’ cats and sympatric domestic cats in the Daursky Reserve (Russia), have been exposed to feline immunodeficiency virus and feline leukemia virus (41). To date, exposure or infection with FCoV-1 or -2 has not been reported for free-ranging or captive Pallas’ cats. However, studying if this species is susceptible to FCoV infection and the development of FIP is essential, given the high prevalence of FCoV in domestic cats worldwide and in other wild felids living in proximity to the known distribution range of the Pallas’ cat (29). Since it is difficult to study free-ranging Pallas’ cat, assessing infectious diseases in animals in captivity is essential to understand this species’ susceptibility to certain viruses (42).

Here, we report a retrospective molecular epidemiological investigation of FCoV infection in a zoological institution in the U.S. in which a Pallas’ cat kitten and a domestic cat that shared the same room succumbed to FIP in 2008. In this study, we used molecular and histological tools to characterize the partial genome sequence of the lethal FCoV detected in both individuals. This study provides the initial genetic description of an FCoV-1 that caused FIP and the eventual death of a wild felid. To our knowledge, we provide the first genetic evidence of FCoV-1 transmission between a captive wild felid and a domestic cat. Our results also highlight the epidemiological relevance of characterizing the FCoV genotypes circulating in wild felids.

## Methodology

### Coronavirus screening

In November 2008 two Pallas’ cat kittens (less than 5 months old) and a female short-haired domestic cat (DCB091) died at a research colony in a zoological institution in the U.S. The lung, pleural tissue, and intestine from one of the Pallas’ cat kittens (OM1164) and the mesenteric lymph node of the domestic cat were collected and frozen at -80 °C until further use. Frozen tissues were thawed on ice and then equilibrated to room temperature. Thawed tissues were formalin-fixed, paraffin-embedded (FFPE) at the Section of Anatomic Pathology, Department of Biomedical Sciences at the Cornell University College of Veterinary Medicine Animal Health Diagnostic Center (AHDC). For each tissue, an unstained slide of 5 μm thick tissue sections were used. A probe directed to the RdRp gene (39) was used to detect FCoV RNA by *in-situ* hybridization (ISH, RNAscope ® Probe V-FIPV-ORF1a1b, Advanced Cell Diagnostics). The ISH process was carried out at the AHDC in the automated staining platform Ventana Discovery Ultra (Roche Tissue Diagnostics) using the Discovery kits for mRNA Sample Prep, mRNA Red Probe Amplification and mRNA Red Detection and the Ventana Hematoxylin and Ventana Bluing Reagents for counterstaining.

Approximately 40 mg of each tissue was used for RNA extraction using the Monarch^®^ Total RNA Miniprep Kit (New England Biolabs, NEB) according to the manufacturer’s instructions and including the suggested DNaseI in-column step. Synthesis of cDNA was carried out using the LunaScript® RT SuperMix Kit (NEB) following the manufacturer’s instructions. Screening for coronavirus was carried out using previously described primers targeting a conserved region of the RNA-dependent RNA polymerase (RdRp) gene of FCoV (35) using the Platinum II Taq Hot-Start DNA Polymerase (Thermo Fisher Scientific). Positive samples were sequenced in the MinION Mk1b (Oxford Nanopore Technologies, ONT) using a Flow Cell R10.4 (ONT) and the Native Barcoding Kit 24 V14 (ONT).

### Feline coronavirus sequencing

Positive samples were further sequenced using a hybridization capture (HC) enrichment followed by next-generation sequencing (NGS). The HC method uses a panel of single-stranded oligonucleotide “baits” to capture by hybridization targeted sequences from cDNA libraries prepared from clinical samples. The “captured” libraries are then sequenced using NGS. To detect and sequence both FCoV-1 and -2 two panels were designed by Twist Biosciences. The first panel contained the sequences of the complete S gene of FCoV-1 Black and FCoV-2 79-1146 (accession numbers EU186072 and DQ010921, respectively). The second panel included the sequences of the complete 3’-end of 141 variants of FCoV-1 (Supplementary Table 1). The cDNA obtained for the initial viral screening was used to synthesize dsDNA using 5U of DNA Polymerase I, Large (Klenow) Fragment (NEB). The resulting dsDNA was used to construct TruSeq-compatible libraries using the Twist Library Preparation EF Kit 2.0 and the Twist CD Index Adapter set 1-96 (Twist Biosciences). For the HC the Twist Hybridization and Wash kit and the Twist Universal Blockers (Twist Biosciences) were used. The HC was carried out for 16 hours. Obtained libraries were sequenced in the iSeq 100 Sequencing System (Illumina) using the iSeq100 i1 Reagent V2 (300-cycles). For the library and HC assays, DNA was quantified in the Qubit 4 Fluorometer (Thermo Fisher Scientific) using the dsDNA High Sensitivity Assay kit (Thermo Fisher Scientific).

### Genetic analysis

Reads were mapped to the 143 sequences of FCoV-1 (Supplementary Table 1) and -2 that were included in the two panels (Supplementary Table 1) using BWA-MEM2 (43, 44) in Galaxy V 22.01 (45). Mapped reads were visualized in Geneious Prime 2023.0 (Dotmatics) and consensus sequences were generated using a 75% threshold. Assembled sequences were uploaded to GenBank (accession numbers PP067758 - 61). The complete 3’-end of the genome (8.2 Kb) of the two variants obtained in this study was aligned with 30 homologous sequences from other FCoV-1 using the MUSCLE algorithm (46) in Geneious Prime 2023.0 (Dotmatics). Gaps were manually deleted from this alignment. The resulting gap-free alignment was used to detect possible recombination breakpoints using the Recombination Detection Program (RDP 5, (47)), using the RDP method (48).

The two partial sequences of the RdRp gene (278 nucleotides, nt) obtained in this study were aligned with 56 other homologous sequences from other members of the *Alphacoronavirus-1* species including FCoV-1, FCoV-2, CCoV-1, CCoV-2, and TGEV. The two sequences obtained from two cheetahs (Aju92 and Aju93) during the 1982/1983 outbreak (accession number EU664236, EU664268) were also included. The partial sequences of the S gene (3,681 nt), and the E (243 nt), M (867 nt), N (1,131 nt), and 7a (324 nt), b (618 nt) genes were each aligned with 93 other homologous sequences. The alignments were obtained using the MUSCLE algorithm (46) in Geneious Prime 2023.0 (Dotmatics). For each alignment, the best-fitting nucleotide substitution model was obtained in MEGA 11 (49). Maximum-likelihood (ML) phylogenetic trees were constructed for each gene using the PhyML 3.0 algorithm (50) and 1,000 bootstraps to test the branch support. Trees were visualized and colors were modified in MEGA 11. Phylogenetic trees could not be constructed for genes 3a, b, and c due to deletions in numerous sequences. Pairwise percentage similarity of the nucleotide and amino-acid sequences of each gene was calculated using Geneious Prime 2023.0 (Dotmatics).

## Results

PCR and ISH targeting the RdRp gene detected FCoV RNA in the lung of the Pallas’ cat kitten and the mesenteric lymph node (MLN) of the domestic cat (Figure 2). FCoV RNA was not detected in the pleural tissue or the intestine of the pallas cat. The sequence of the partial region of the RdRp gene (278 nt) of the Pallas’ cat kitten (OM1164) and the domestic cat (DCB091) was 99.6% similar (only one nucleotide difference between the sequences). Phylogenetic analysis of the partial region of the RdRp sequenced revealed that the sequences are within the FCoV-1/FCoV-2 group (Figure 3), and group with an FCoV-1 variant obtained from a domestic cat in the U.S. in 2002 (FCoV-1 RM USA 2002) and the two variants obtained from two cheetahs during the 1982 Oregon case (AJU92 and AJU93) from which the genotype is unknown. Genetic comparison of the RdRp sequence obtained from the Pallas’ cat and the other wild felids showed 95% similarity with the cheetah sequences and 91.8% with the sand cat sequence.

**Figure 2.**
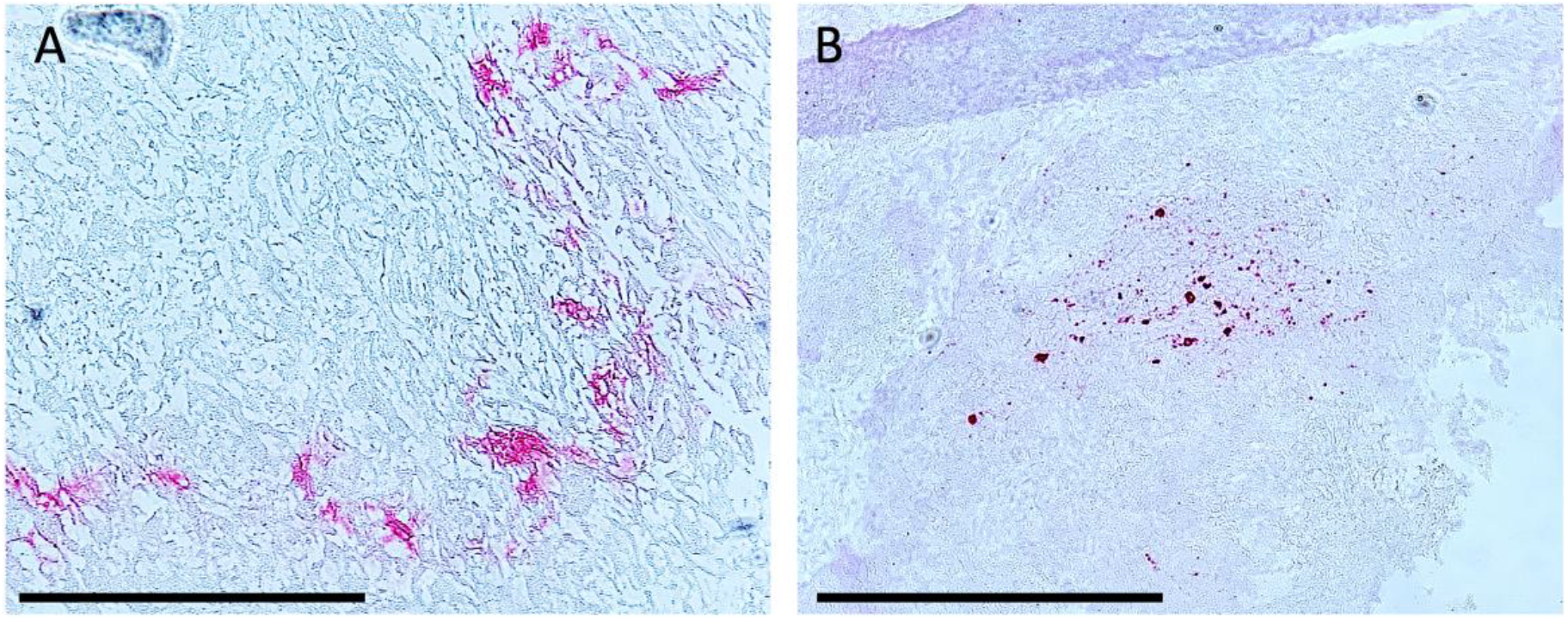
Detection of FCoV RNA (in magenta) in tissues by *in-situ* hybridization. A. Lung of the Pallas’ cat kitten. B. Mesenteric lymph node of a domestic cat. The scale bar indicates 360 μm.

**Figure 3.**
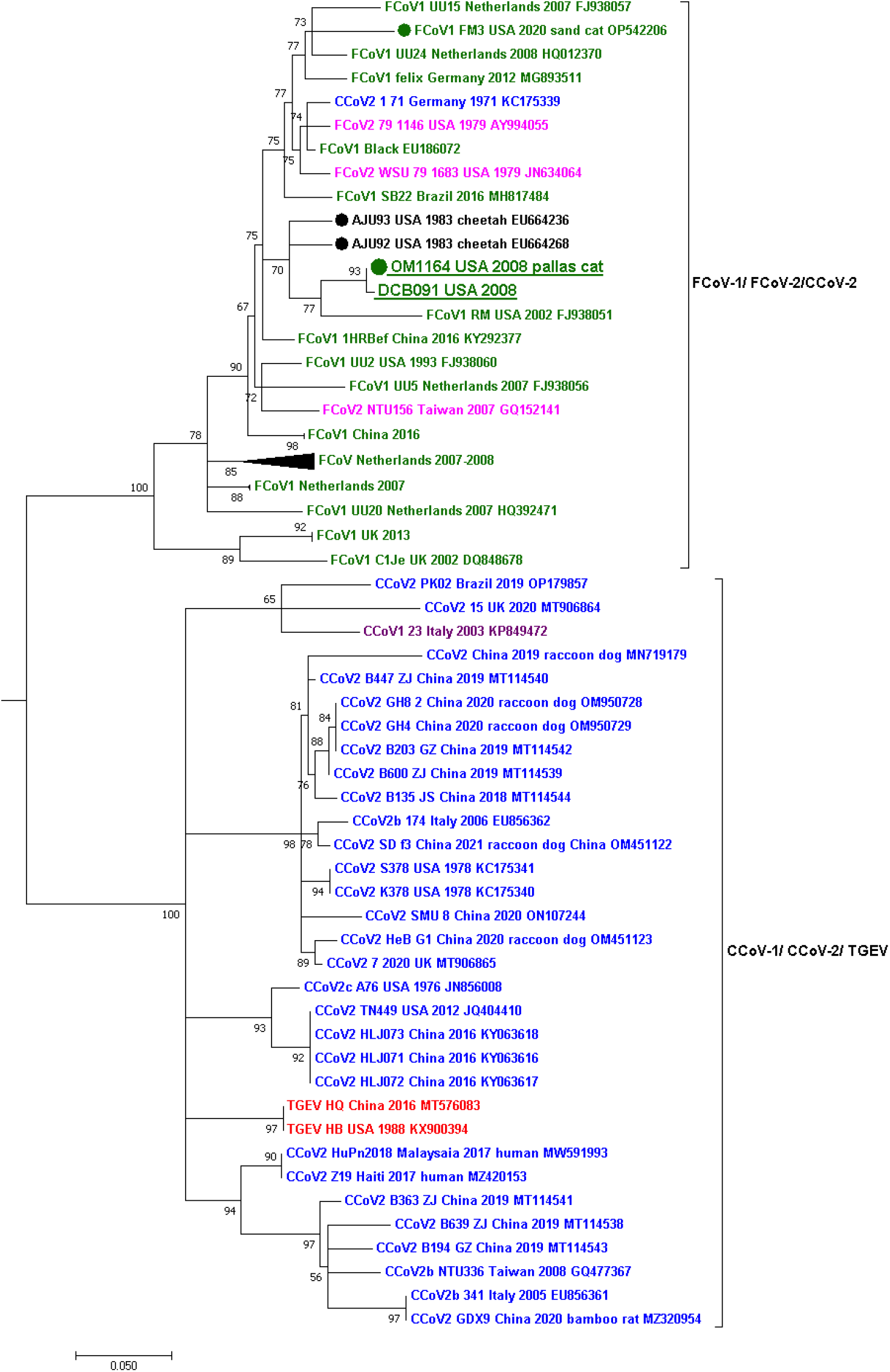
Maximum likelihood phylogenetic tree of a partial region of the RdRp gene (278nt) of the *Alphacoronavirus-1* species. Each genotype is shown in a different color: FCoV-1 in dark green, FCoV-2 in fuchsia, CCoV-1 in purple, CCoV-2 in blue, and TGEV in red. Two groups are observed in the tree, one grouping FCoV-1, FCoV-2, and CCoV-2 and another grouping CCoV-1, CCoV-2, and TGEV. The sequences obtained in this study from the Pallas’ cat (OM1164) and the domestic cat (DCB091) are underlined and placed in the FCoV-1/FCoV-2 group. Within this group, the four sequences obtained from wild felids are marked with a dot accompanied by the host species’ common name. The two FCoV of unknown genotype obtained from two cheetahs from the 1982 Oregon case are in black. For all viruses, the virus name, year and country of collection, and NCBI GenBank accession numbers are shown at the tip of the tree. Numbers at the branches indicate bootstrap percentage values from 1000 replicates. Groups of identical or very similar sequences were grouped and are shown as triangles. Triangle size is correlated with the number of sequences in the group. Branches with support <50 were collapsed. Nucleotide substitution model used: HKY+I+G.

The sequence of the 3’-end of the genome (8.2Kb) was obtained for the FCoV of both the domestic cat (DCB091) and the Pallas’s cat kitten (OM1164). This genome region includes a partial sequence of the S gene (3,681 nt long, missing around 600 nt of the 5’-end of the S1 domain), and genes 3a, 3b, 3c, E, M, N, 7a, and 7b (Figure 2). Genetic comparison of the nucleotide sequence of the whole 3’-end revealed that the two variants share 98.1% pairwise nucleotide similarity. Genetic comparison of the amino-acid sequence of the structural and accessory proteins revealed that the S and 7b proteins were the most variable (97.2% pairwise amino-acid identity), and only the 3a gene was identical between the two viruses (Figure 4). Amino-acid sequence analysis of the S protein revealed an FCS in the S1/S2 region in both variants, indicating that both are FCoV-1 (Figure 4). The FCS of both variants differs by one residue in position P1, which is an S in the domestic cat and an R in the Pallas’ cat (Figure 4). In position P3, both variants have an A (Figure 4). Both variants had the same sequence in the S2’ cleavage site (GKRS), but there was a region of high variability right before the S2’ cleavage site (Figure 4). In site ‘1058’, both variants had L, and no deletions or insertions were detected in genes 3c, 7a, or 7b (Figure 4). Phylogenetic analysis of the partial region of the S gene revealed that the two sequences group together and are within the FCoV-1 group (Figure 5). These two sequences are within a group with an FCoV-1 obtained from a cat in the Netherlands in 2008 (UU24) and with the sequence from the sand cat obtained in 2020 (FM3, Figure 5). No reads mapped against the S gene of FCoV-2, showing that there was no co-infection with the two genotypes. Phylogenetic trees using the other structural genes, including E, M, N, and 7ab, show that the two variants obtained in this study always group together and are within the FCoV-1/FCoV-2 group (Supplementary Figure 1). No recombination breaking points were detected in the two variants obtained in this study.

**Figure 4.**
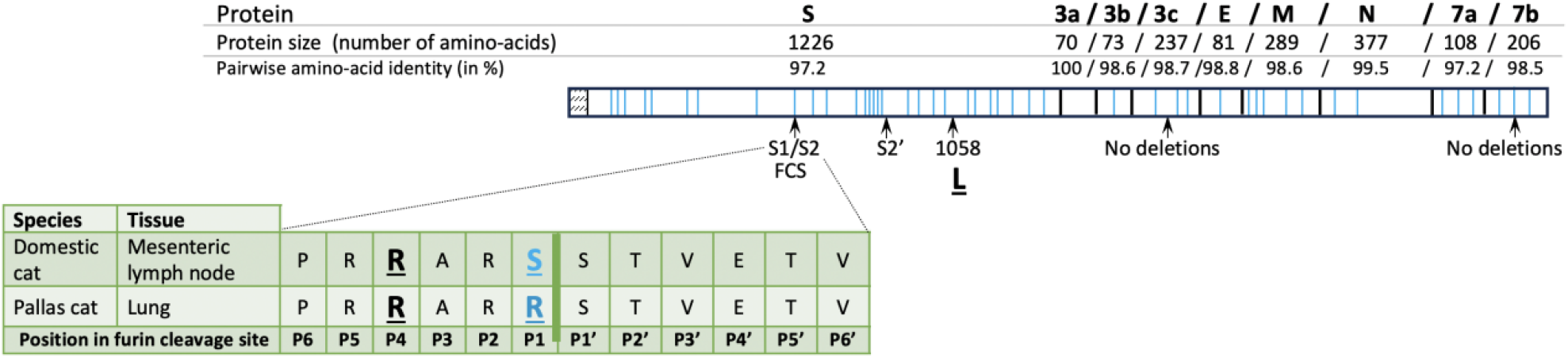
Schematic representation of the 3’-end of the genomes (∼8.2Kb) of the FCoV-1 detected in the domestic cat and the Pallas’ cat. The name of the proteins, size, and percentage amino acid identity is shown above the genome scheme. Light blue vertical lines within each protein indicate the position of amino acids that vary between the two variants. Below the 3’-end diagram pointed with arrows are genome regions relevant for FCoV-1 pathogenicity. The amino acid sequence of one of these regions (S1/S2) is zoomed in to show the furin cleavage sites (FCS) of both variants. Amino acids that vary between the two sequences are highlighted in light blue. Residues previously reported to be related to pathogenicity are underlined and include positions P4 and P1 in the FCS and residue 1058.

**Figure 5.**
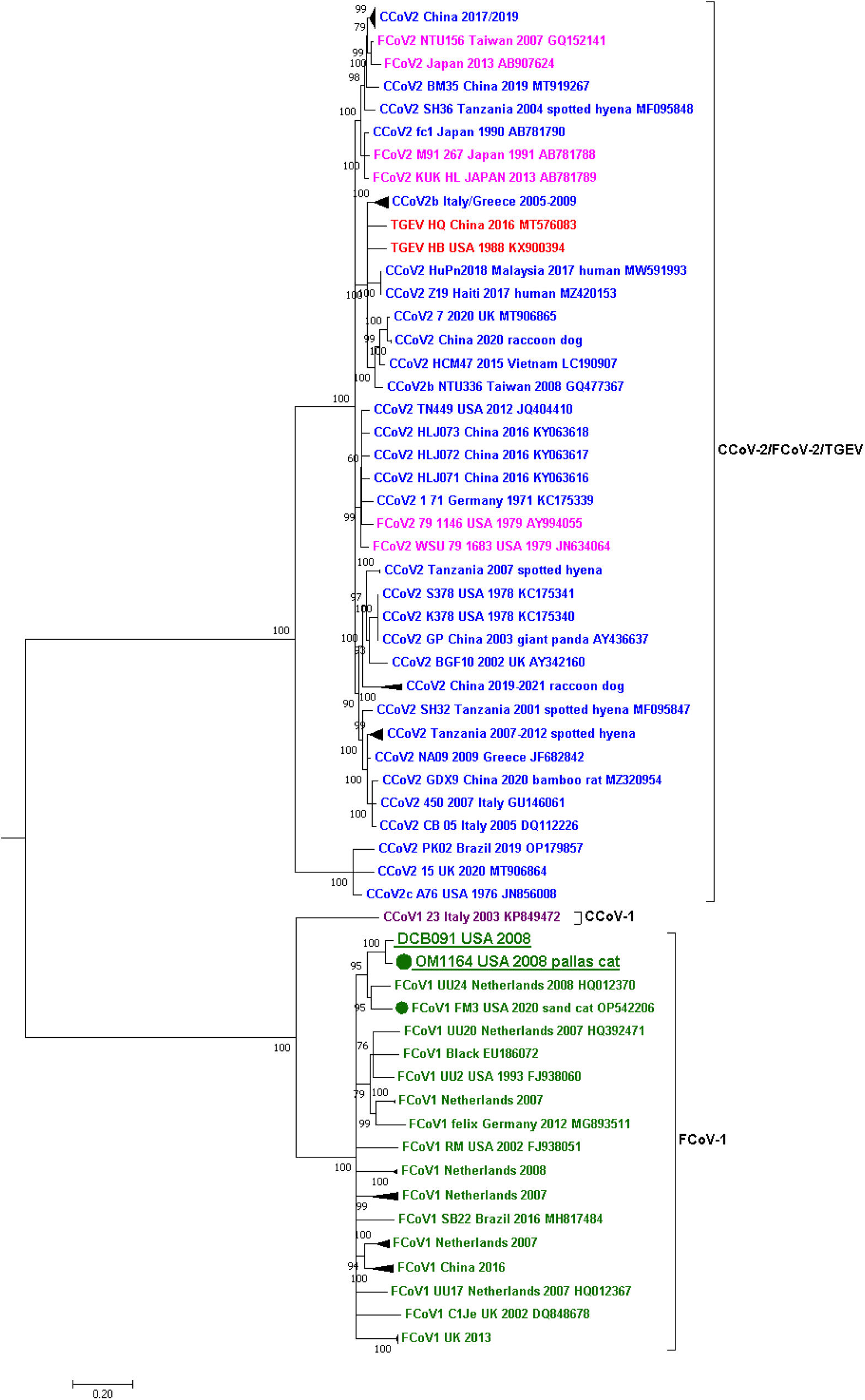
Maximum likelihood phylogenetic tree of a partial region of the S gene (3,681 nt) of the *Alphacoronavirus-1* species. Each genotype is shown in a different color: FCoV-1 in dark green, FCoV-2 in fuchsia, CCoV-1 in purple, CCoV-2 in blue, and TGEV in red. Two groups are observed in the tree, one grouping all variants of FCoV-1 and another grouping FCoV-2, CCoV-2, and TGEV. CCoV-1 is outside these groups. The sequences obtained in this study from the Pallas’ cat (OM1164) and the domestic cat (DCB091) are underlined and placed in the FCoV-1 group. Within this group, the two sequences obtained from wild felids are marked with a dot accompanied by the host species’ common name. For all viruses, the virus name, year and country of collection, and GenBank accession numbers are shown at the tips of the tree. Numbers at the branches indicate bootstrap percentage values from 1000 replicates. Branches with support <50 were collapsed. Groups of identical or very similar sequences were grouped and are shown as triangles. Triangle size is correlated with the number of sequences in the group. Nucleotide substitution model used: HKY+I+G. A total of 95 sequences were included in the analysis.

## Discussion

In this study, by using next-generation sequencing techniques and *in-situ* hybridization we detected and characterized FCoV-1 in a Pallas’ cat and a domestic cat that shared the same room at a zoological institute in the U.S. (Figure 1, 2). The major difference between FCoV-1 and FCoV-2 is their S gene because FCoV-2 is a recombinant genotype that acquired its S gene from CCoV-2 (Figure 1). Therefore, to determine the genotype of FCoV, sequencing of either the complete S gene or a part of it is required. When using the genes commonly used for detecting FCoV in wild felids, including RdRp (Figure 3) and 7ab (Supplementary Figure 1), two groups of *Alphacoronavirus-1* can be clearly distinguished. One group includes FCoV-1, FCoV-2, and one strain of CCoV-2 (1-71), while the other group includes CCoV-1, CCoV-2, and TGEV (Figure 3). In the phylogeny of the RdRp gene, the two sequences obtained in this study were placed together within the FCoV-1/FCoV-2 group. Within this group were also the two FCoVs obtained from two cheetahs from the 1982 Oregon (Figure 2). Although there are only a limited number of FCoV-2 sequences available (in fuchsia in Figure 3), they do not form a monophyletic group in the RdRp gene phylogeny (Figure 3). Therefore, when only using the RdRp gene, like in the case of the two cheetah sequences, it is possible to determine if the detected virus is an FCoV (*i*.*e*., if it groups within the FCoV-1/FCoV-2 group), but it is not possible to confirm the specific FCoV genotype (*i*.*e*., if it is an FCoV-1 or an FCoV-2). This is also true for the other structural and accessory genes like E, M, N, and 7ab (Supplementary Figure 1). In contrast, the phylogeny of the partial S gene (Figure 5), revealed two different groups. One includes FCoV-1, and the other includes FCoV-2, CCoV-2, and TGEV (Figure 5). Another characteristic differentiating FCoV-1 and -2 is that the S protein of FCoV-1 has an identifiable FCS in the S1/S2 region. The two FCoV sequences obtained in this study were placed within the FCoV-1 cluster (Figure 5), and their S protein had an FCS in the S1/S2 region (Figure 4). These results confirm that the genotype infecting both the Pallas’ cat and the domestic cat is an FCoV-1. Although clinical manifestations consistent with FIP were previously reported in one Pallas’ cat individual (51), no genetic or histological evidence of FCoV or FIP had been previously reported for this species. Therefore, this study provides the first genetic evidence that FCoV-1 can cause FIP in the Pallas’ cat.

The high mortality associated with FIP in the cheetah population at an Oregon safari park, in which 60% of the cheetah population died, highlights the need to understand the risk factors involved in the development of FIP in wild felids. This knowledge is crucial for understanding why FIP has only been found in captive cheetahs, despite FCoV being detected in both free-ranging and captive cheetahs. One hypothesis is that the accumulation of certain mutations in FCoV, may allow mutant variants to acquire macrophage tropism, a critical step for the development of FIP (5). This is known as the ‘internal mutations’ hypothesis (4). A recent study found that mutations in specific residues of the S protein of FCoV-1 and FCoV-2 were strongly associated with FIPV (19). No such association was found in 3abc or 7b (19), proteins previously thought to be involved in the development of FIP (22). Therefore, detecting the specific genotype causing FIP in wild felids and studying its S protein is crucial to determining the molecular mechanisms involved in the emergence of pathogenic variants in wild felids. For FCoV-1, mutations that disrupt the FCS have been reported in most FIP cases (12). In one of the positions within the FCS associated with FIPV (P4), we did not detect mutations related to FIPV (both sequences had an R in P4, Figure 4). We detected a mutation in residue P1 of the FCS of the FCoV-1 detected in the mesenteric lymph node of the domestic cat (R→S, Figure 4) but not in the one detected in the lung of the Pallas’s cat. This mutation in P1 was also identified in the FCoV-1 detected in the kidney of a sand cat (32). Mutations in this position abrogate furin cleavage and are associated with FIPV (12). Previous reports have shown that different mutations in the FCS of FCoV-1 can be detected in different tissues of cats with FIP (52). As only the mesenteric lymph node of the domestic cat and the lung of the Pallas’ cat were available in this study, it is only possible to suggest that the difference observed in the FCS is due to its detection in different tissues. In contrast, both variants had the residue L at position 1058, which is considered a marker for FIPV (20) (19). This example shows that detecting mutations related to FIP is complex and requires studying various regions in S and multiple tissues in affected individuals.

Sequencing of the 3’-end of the genome showed that the FCoV from the Pallas’ cat and the domestic cat were highly similar (98.1% pairwise nucleotide similarity). Also, the two always grouped together in the phylogenetic trees of the RdRp and all the structural genes, including the S gene (Figures 2 and 3, Supplementary Figure 1). A comparison between the two sequences of the 3’-end of the genome revealed that most of the mutations detected between the FCoV-1 from the domestic cat and the one from the Pallas’ cat are in the S gene, while the other genes were highly conserved (>98.5% pairwise sequence similarity), including gene 3a that was identical between the two FCoV-1 (Figure 4). This suggests that the two viruses have the same origin, but most mutations occurred in S because this gene is involved in pathogenicity and tissue tropism. The observed mutations in S can also be related to host switching, as has been observed for another *Alphacoronavirus-1* (53). Overall, these results support the idea of the “internal mutation” hypothesis (4) and are the first genetic evidence of the cross-species transmission of FCoV-1 between a wild felid, the Pallas’ cat, and a domestic cat. It is essential to understand that FCoV-1 can be transmitted between domestic and wild felids to develop mitigation strategies that diminish the intraspecific spread of this virus. This is particularly important in cities where free-roaming feral or stray cats are highly abundant and pose a major public health risk due to their potential contact with urban wildlife and humans. FCoV has been reported in urban (54) and rural feral/stray cats (55) as well as in household cats that are allowed to go outdoors (56). In the U.S., it is common to feed feral/stray cats, which often leads to the formation of cat colonies near these areas. The cat food can also attract local wildlife, which may include wild felids. Many studies have reported that FCoV prevalence is higher in environments with multiple cats, including multi-cat households (57) and rescue centers (58). Therefore, urban areas where human-provided food promotes the grouping of feral/stray cats and wild felids may increase the risk of cross-species transmission of FCoV. An ongoing outbreak of a novel FCoV (FCoV-23) in feral/stray cats and free-roaming owned cats in Cyprus has resulted in at least 91 confirmed cases since 2021, representing a 40-fold increase in reported cases of FCoV-related deaths in the island. (59). This outbreak highlights the negative impact FCoV could have on populations of both domestic and wild felids and the need for rapid genetic characterization of FCoV.

Co-infection with both genotypes of FcoV has been detected in several domestic cats diagnosed with FIP (7). However, the prevalence of co-infection in FIP cases has not been thoroughly assessed in domestic or wild felids. Studying the co-infection of FCoV-1 and FCoV-2 in FIP cases is essential to evaluate the contribution of each genotype to the development of the disease and to determine if there is a synergistic effect when co-infection occurs. The method presented in this study included baits to detect by hybridization capture (HC) and sequence using NGS FCoV-1 and FCoV-2 simultaneously. Therefore, it is essential to implement HC and NGS to carry out epidemiological surveillance of both FCoV genotypes in domestic and wild felids. Additionally, FCoV-2 with different recombination breaking points have been reported in domestic cats, suggesting that multiple recombination events between FCoV-1 and CCoV-2 have occurred (7). Studying and genetically characterizing FCoV genotypes in wild felids (captive or free-ranging) will help determine if multiple recombinant variants of FCoV-2 circulate in different wild species. Overall, to determine the origin of recombinant FCoV-2 genotypes in domestic and wild felids, it is important to study the prevalence not only of FCoV-1 and FCoV-2, but also of CCoV-2. For example, CCoV-2 has been reported in several wild species of carnivores, including feliforms like the spotted hyena *(Crocuta crocuta*, (53)). However, knowledge of natural infection of CCoV-2 in domestic or wild felids remains limited. To study co-infection, sequencing PCR products, using Sanger sequencing or even NGS techniques like Oxford Nanopore Technologies, may not be sufficient as using primers biases the detection of certain genotypes. This is especially true when targeting the S gene of *Alphacoronavirus-1*, which does not always yield reliable results (60). In contrast, it has been shown that by using HC, it is possible to detect and sequence complete genomes of multiple CoVs in a single sample, as well as to discover new CoVs (61). Implementing the hybridization capture method reported in this study to characterize different *Alphacoronavirus*-1 and to identify co-infections will be essential to determine if wild felid species could act as mixing vessels for the emergence of recombinant genotypes of FCoV or *Alphacoronavirus-1* in general.

## Acknowledgements

Work in the Whittaker lab is funded in part by the Cornell Feline Health Center. We thank William F. Swanson and Meredith Brown for collecting and sending the samples to Cornell.

