## Supplemental Table 1 and Supplementary Figure 1 for "Molecular detection using hybridization capture and next-generation sequencing reveals cross-species transmission of feline coronavirus type-1 between a domestic cat and a captive wild felid": FCoV_pallas_cat_supplementary_info.docx

Supplementary Table 1. Accession numbers of the 141 sequences of FCoV-1 included in the second panel to detect and sequence FCoV.

Accession number

FJ938053

KP143509

KY566211

FJ938059

HQ012371

GU553362

KP143508

HQ392470

EU186072

HQ012368

KU215427

AB088222

GU553361

HQ392471

KX722529

HQ012370

MW316834

MW316840

MT444152

KJ665876

MN165107

FJ917530

FJ917520

JN183882

FJ917519

FJ938059

HQ012371

AB695067

FJ938051

FJ938053

MW316839

KY292377

FJ917531

MW316841

FJ917522

FJ938055

FJ917524

FJ938052

MW316837

FJ917535

MW316836

MW316845

KX722531

KY566209

MF457591

MW030110

MW316844

MW316830

MW316833

MW030108

MW316846

KF530123

D32044

KY566211

FJ938054

KJ665866

JN183883

MW316831

DQ160294

KX722530

MW316832

KJ665862

MW316838

HQ012372

FJ917523

MW316847

HQ392469

KY566210

FJ917534

FJ917521

MT239440

FJ938062

MG893511

AB535528

MW316835

FJ938056

MW316842

MW316843

FJ938058

MH817484

HQ392472

EU186072

KU215421

AB088222

HQ392471

FJ917520

HQ012371

HQ392469

HQ012372

FJ938058

MF457591

HQ012368

FJ938053

FJ938056

MW316837

KP143509

MW316835

MW316846

KY292377

MW316840

MW030108

MN165107

MW316833

MW316847

AB535528

MW316834

KX722531

FJ938055

MW316836

MW316830

MW316831

MW316845

MW030110

KF530123

MW316839

JN183882

MW316832

D32044

FJ938052

MT444152

KY566210

DQ160294

HQ012367

MW316844

DQ848678

KX722529

MW316842

HQ012370

MW316843

HQ392470

MW316841

MW316838

MT239440

FJ938051

FJ938062

MG893511

KY566211

FJ938059

AB695067

KX722530

KY566209

Supplementary Figure 1A Phylogenetic tree of the E gene.

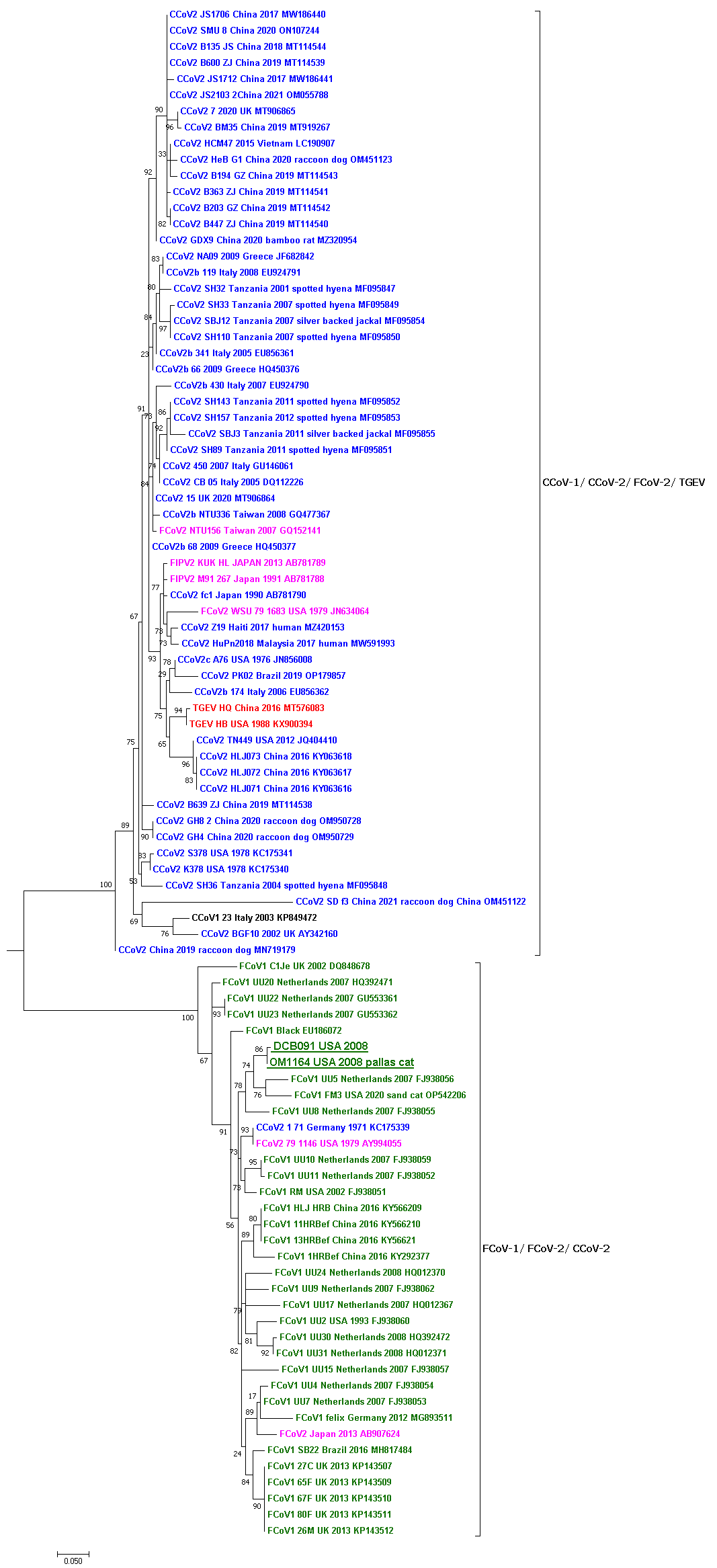

Supplementary Figure 1B Phylogenetic tree of the M gene.

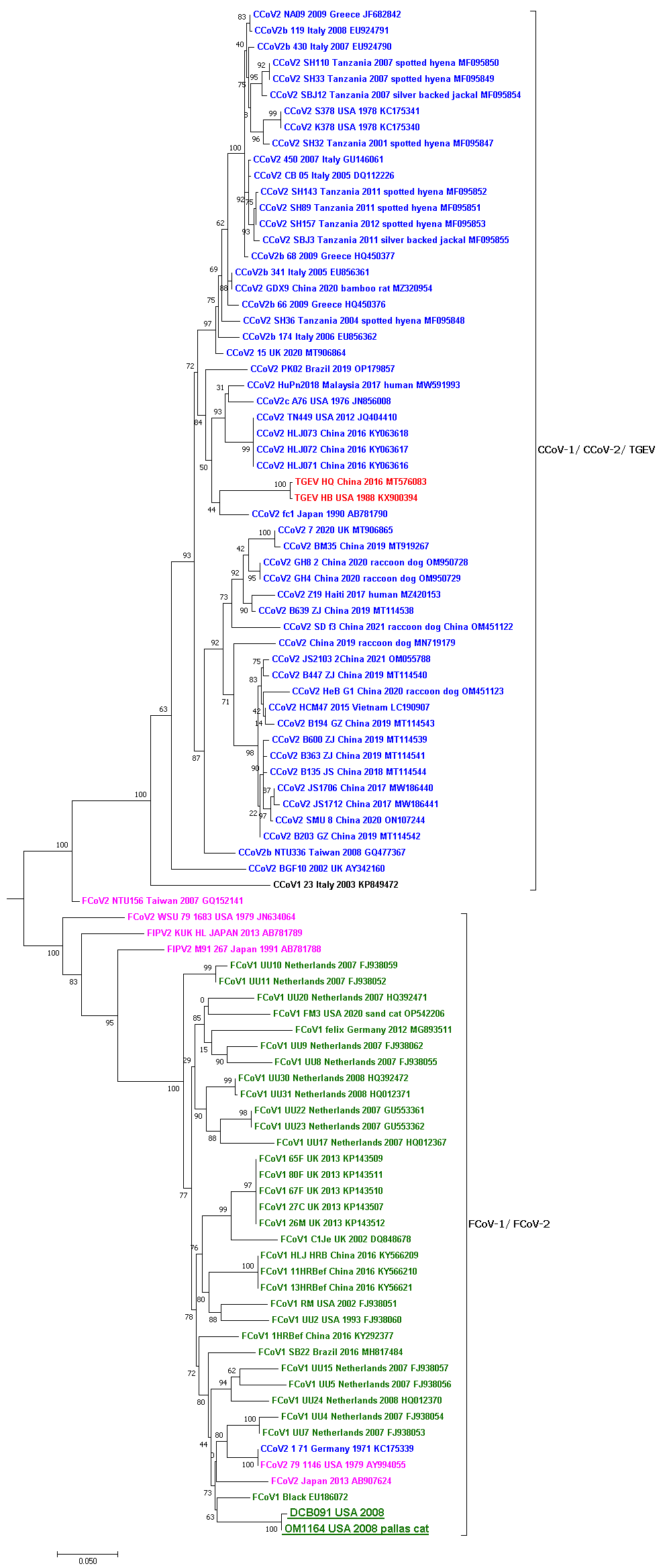
Supplementary Figure 1C Phylogenetic tree of the N gene.

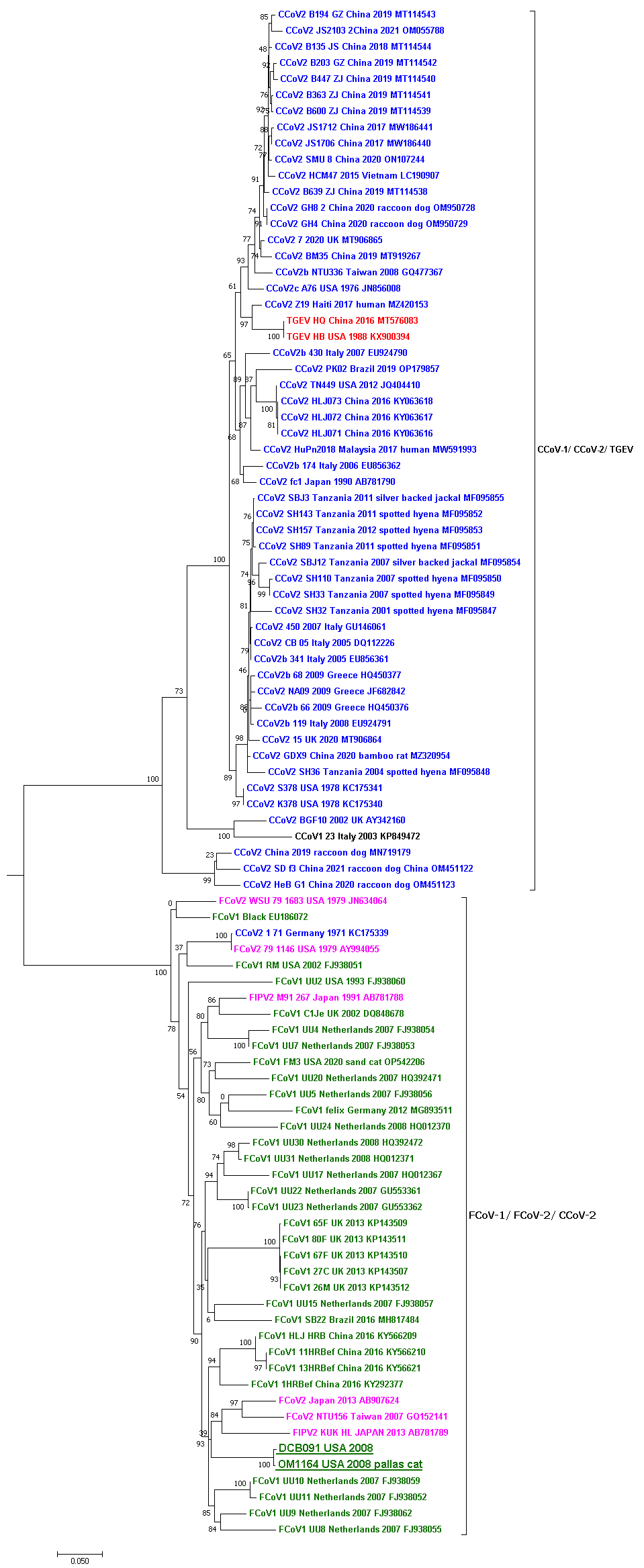

Supplementary Figure 1D Phylogenetic tree of the 7ab genes.

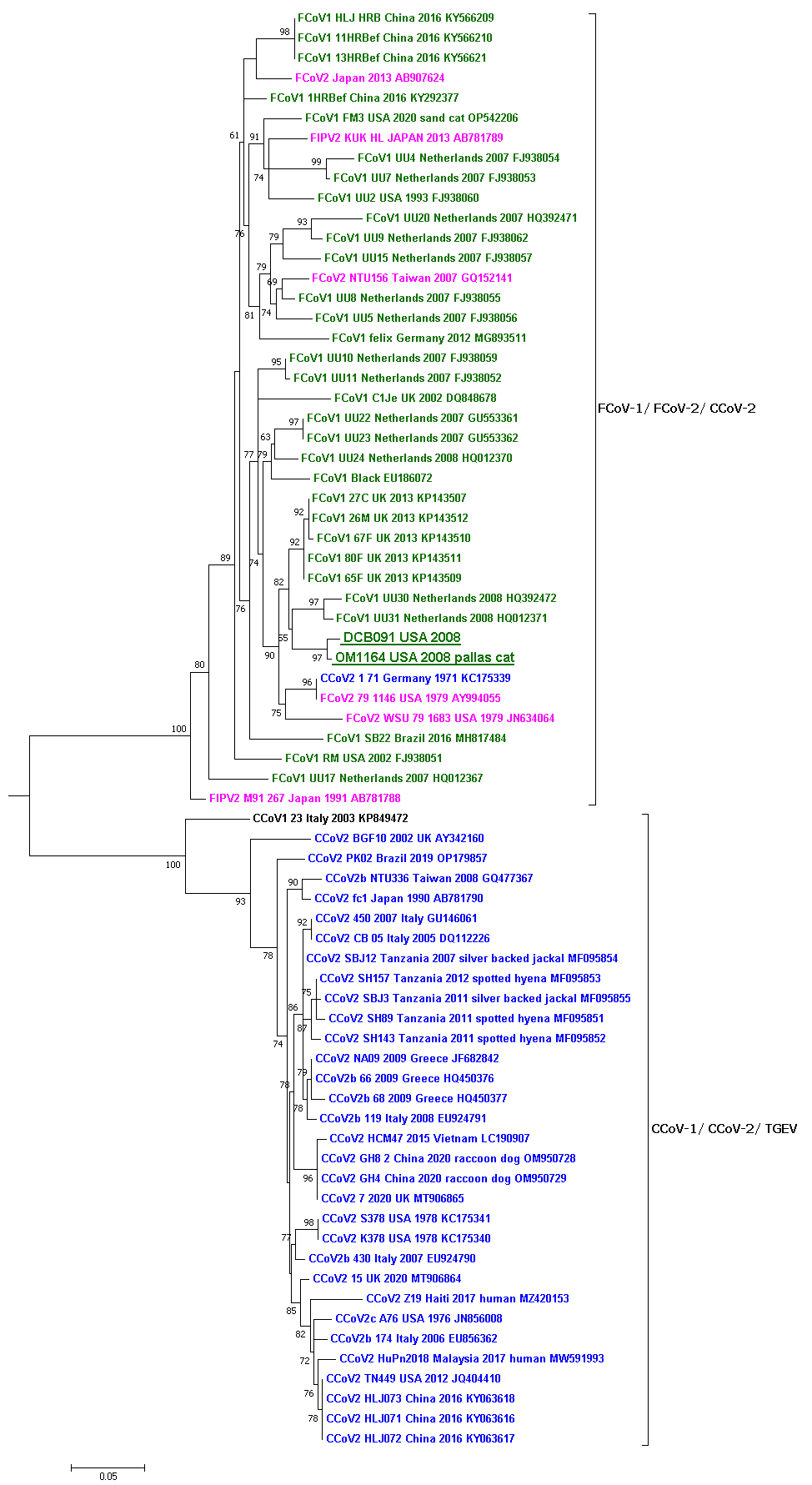
